# Quantitative Real-Time PCR Detection of Inactivated H5 Avian Influenza Virus in Raw Milk Samples by Miniaturized Instruments Designed for On-Site Testing

**DOI:** 10.1101/2025.06.02.657307

**Authors:** Chih-Chi Hsiao, Chiu-Chiao Lin, Mike Li, Patrick Lin, Ya-Mei Chen, Ming-Chu Cheng, Sherrill Davison, Jianqiang Ma, Hai-Lung Dai

## Abstract

Highly Pathogenic Avian Influenza Virus of H5 (HPAI A(H5N1)) subtype has emerged as one of the most important zoonotic pathogens with significant economic consequences. The recent outbreak of H5N1 avian influenza in dairy cattle in the United States highlights the importance of early detection in managing and mitigating HPAI A(H5N1) outbreaks. A new diagnostic platform (the MT platform) consisting of miniaturized instruments for DNA/RNA purification and RT-rtPCR, designed for mobilse, on-site testing, is compared with a platform of benchtop instruments (QIAGEN RNeasy and QuantStudio™5) for detecting inactivated HPAI A(H5N1) virus spiked into raw milk samples. Results show that, despite the presence of inhibitors in raw milk, HPAI A(H5N1) virus can be detected in all samples using both platforms. The MT platform shows higher sensitivity than the benchtop platform: the MT Ct values are ∼2 units lower than the benchtop Ct values. Our findings demonstrate the robustness of the MT platform for detecting HPAI A(H5N1) virus in raw milk samples and support its use as an on-site detection and screening for rapid surveillance and response.

## INTRODUCTION

Avian Influenza Virus has been a significant threat to the poultry industry worldwide, with global outbreaks primarily caused by the H5 and H7 subtypes since the mid-20^th^ century[1]. The current wave of avian influenza began in 2020 and is dominated by the highly pathogenic avian influenza virus H5 subtype (HPAI A(H5N1)) strain of clade 2.3.4.4b, which originated in Europe and first spread to North America in 2021, with frequent spillover events to mammals reported[1,2]. In the United States, the potential spread of the HPAI A(H5N1) strain beyond poultry increased significantly after HPAI A(H5N1) was isolated from milk in dairy cattle in Texas in February 2024, with cattle showing symptoms such as reduced milk production and decreased feed intake[3]. Zoonotic infection of HPAI A(H5N1) virus of dairy farm workers has also been reported, although the symptoms were milder compared to bird-transmitted HPAI A(H5N1) infection[4,5].

The implications in human public health have been further highlighted by the fact that 70 confirmed zoonotic HPAI A(H5N1) human cases have been reported in the United States as of May 2025 [6,7]. A new genotype D1 typically carried by wild birds was found to be responsible for the first severe HPAI A(H5N1) case in Canada in November 2024, and the first severe case in the United States in Louisiana in December 2024 which resulted in death [8,9].

Since its first discovery in cattle milk, the HPAI A(H5N1) virus has spread in the United States. As of May 2025, at least 1,052 dairy herds across 17 states have been affected, prompting urgent governmental action to address the growing crisis (U.S. Centers for Disease Control and Prevention, 2025). California, the largest milk-producing state in the United States, bears the brunt of this outbreak, accounting for 73% of cases reported in the United States [6]. In response, the state government of California has declared a state of emergency to mobilize resources and combat the virus’s rapid spread, while upholding consumer confidence in dairy products (State of California, 2024). Nationwide, to address the importance of early detection and containment to mitigate the spread of HPAI A(H5N1) virus within dairy herds and protect public health, the United States Department of Agriculture (USDA) announced a new Federal Order to implement a National Milk Testing Strategy on December 6, 2024 [11]. Per the Order, milk samples were to be tested by molecular testing using reverse transcription real time polymerase chain reaction (RT-rtPCR), which is considered the standard method used in the detection of HPAI A(H5N1) virus according to the Animal and Plant Health Inspection Service of the USDA [12].

The implementation of the Federal Order for milk testing will challenge the current capacity of RT-rtPCR diagnostic testing in many state diagnostic laboratories. Further, the current laboratory diagnostic routine from taking samples to transportation to a laboratory to results takes time that is critical for tracing the original source of infection. Here we present a new model of RT-rtPCR detection, namely the Mobile Testing (MT), using miniaturized extraction^a^ and PCR instruments^b^ that can be housed in a mobile laboratory for conducting the diagnostic testing on-site. The MT platform for detecting HPAI A(H5N1) virus in milk samples offers a critical solution for rapid surveillance and early intervention. By enabling on-site and timely detection, the MT platform reduces the lag between sample collection and diagnosis, enhancing the ability to contain outbreaks before they escalate.

This study aimed to evaluate the effectiveness of the MT platform for detecting the HPAI A(H5N1) virus in raw milk samples and its adaptability to different operational settings by comparing different systems.

## MATERIALS AND METHODS

### Virus

Three samples of lysis buffer-inactivated HPAI A(H5N1) (clade 2.3.4.4b) taken from a goose autopsy tissue/organ sample in Pingtung County, Taiwan, were obtained as part of a research grant awarded to Ya-Mei Chen from the Animal and Plant Health Inspection Agency, Taiwan Ministry of Agriculture (research grant: Research and Analysis of Key Avian Disease Surveillance and Control, ID: 113AS-5.5.5-VP-01; In Traditional Chinese: 家禽重要疾病監測及防控研析, ID: 113 農科-5.5.5-檢-01). The concentrations of inactivated virus suspended in the lysis buffer of the three samples are estimated based on quantitative PCR measurements using the positive control of the ThermoFisher VetMAX™-Gold AIV RNA test kit (Cat# 4485261, ThermoFisher Scientific, Waltham, MA) with a known RNA concentration of 1,000 copies/µL. The estimated concentrations of samples 1, 2, and 3 are 1,000, 10,000, and 1,000 copies/µL, respectively. All experiments were conducted in the laboratories of the Schweitzer Biotech Company in Taipei, Taiwan.

### Raw Milk Samples

Fifteen fresh raw milk samples were collected from different dairy farms in Taiwan. All milk samples were diluted with phosphate-buffered saline (PBS) at a ratio of 1:3 by volume to reduce the inhibitory effects of components in the milk. The 15 raw milk samples were used to evaluate the performance of the MT platform, which consists of a DNA/RNA purification instrument (namely “EZextrator”)^a^ and a rtPCR instrument (namely “MiniQ”)^b^ and the USDA-recommended benchtop platform (Qiagen RNeasy + QuantStudio™5) in detecting HPAI A(H5N1) virus. Each of the 15 raw milk samples was spiked with either of the three inactivated HPAI A(H5N1) virus samples (samples 1, 2, or 3) and diluted to reach a concentration of 10 copies/µL. The spiking and dilution process of the raw milk samples is outlined in **Table 1**.

**Table 1.** Inactivated HPAI A(H5N1) virus in milk samples used in this study. First dilution was made by adding 40 µL of inactivated virus sample to 100 µL of milk and 260 µL PBS to achieve a concentration of 25% milk by volume. Subsequent ten-fold dilutions were made by adding 100 µL of the prior sample solution to 900 µL of diluent. *PBS: Phosphate Buffer Solution.

|  | **Sample 1** | **Sample 2** | **Sample 3** |
| --- | --- | --- | --- |
| **Stock concentration** | 1,000 copies/μL | 10,000 copies/μL | 1,000 copies/μL |
| **Diluent** | Milk-PBS* mixture (1:3) | Milk-PBS mixture (1:3) | Milk-PBS mixture (1:3) |
| **Dilution method\*** | Two ten-fold dilutions | Three ten-fold dilutions | Two ten-fold dilutions |
| **Target concentration** | 10 copies/μL | 10 copies/μL | 10 copies/μL |

### Polymerase Chain Reaction

The EZextractor Viral DNA/RNA Extraction Kit (Ref: ATXZ008, Schweitzer Biotech Company, Taipei, Taiwan)^c^ was used to isolate viral RNA using the EZextractor^a^ following the manufacturer’s built-in program. The EZextractor system is a fully automated magnetic-bead-based platform designed to streamline nucleic acid extraction for laboratories handling medium-to large-scale sample volumes. It operates within a compact footprint and features a touchscreen interface for user control. The system supports flexible batch processing with capacities of up to 32 samples per run, with an approximate runtime of 20 minutes. As a comparison in this study, QIAGEN RNeasy Mini kit (Cat. #74106 250, QIAGEN N.V., Venlo, The Netherlands) was used to extract HPAI A(H5N1) virus RNA spiked into raw milk samples using the “Purification of Total RNA from Animal Cells Using Spin Technology” protocol according to the kit handbook.

Extracted HPAI A(H5N1) virus RNA was then amplified by VetMAX™ Gold AIV Detection Kit (Cat# 4485261, ThermoFisher Scientific, Waltham, MA) using Influenza Virus Primer Probe Mix included in the kit (FAM reporter dye). The following control samples were also included in the kit: Influenza Virus-Xeno™ RNA Control Mix for positive control; Xeno™ RNA Control for internal positive control (VIC reporter dye). The cutoff threshold for amplification curves in each PCR run is set according to the VetMAX™ Gold AIV Detection Kit’s user manual, in which, the threshold is set at the 5% of the maximum intensity of the positive control amplification signal level.

The RT-rtPCR process was performed and analyzed using two types of rtPCR thermocyclers: the benchtop QuantStudio™ 5 (QS5) Real-Time PCR System (Cat# A28133, ThermoFisher Scientific, Waltham, MA) and the portable EZcycler Mini real-time PCR (MiniQ) System^b^. The MiniQ PCR System is a compact, space-efficient system for precise quantitative analysis suitable for mobile testing. Equipped with a built-in touchscreen, it operates independently of a computer and has a sample capacity of 16 and three detection channels. Two independent rtPCR runs were performed for each of the milk samples.

The thermal cycling conditions in reverse transcription followed by rtPCR amplification for both thermocyclers are shown in **Table 2**.

**Table 2.** Reverse transcription and rtPCR thermocycling conditions. *: Fluorescence signal is collected at this step.

| Stage | Cycles. | Temperature | Time |
| --- | --- | --- | --- |
| Reverse transcription | 1 | 48°C | 10 minutes |
| RT inactivation/initial denaturation | 1 | 95°C | 10 minutes |
| Amplification: Denaturation | 40 | 95°C | 15 seconds |
| Amplification: Annealing and extension |  | 60°C | 45 seconds* |

### Sensitivity of PCR Detection

Serial ten-fold dilutions of the inactivated virus sample 2 stock solution (10,000 copies/µL) using raw milk-PBS mixture (raw milk samples from farm 8 or farm 9) were performed to determine the limits of detection and the efficiency of the MT platform.

### Raw Milk Inhibitors

To assess whether potential inhibitors in raw milk influence the detection of HPAI A(H5N1) virus, PBS solutions were spiked with inactivated HPAI A(H5N1) virus. Two raw milk samples were randomly chosen from the stocks and diluted with PBS at 1:3 ratio. Each sample, either raw milk in PBS or PBS only, was spiked using the same source with an unspecified quantity of inactivated viruses. Internal positive control was also used to spike the PBS as well as the raw milk diluted with PBS. In this comparison, one set of experiments was performed using the combination of an Ezextractor and the benchtop QS5, and another Ezextractor and the MiniQ.

## RESULTS

Both platforms successfully detected HPAI A(H5N1) virus in all 15 spiked raw milk samples at a viral RNA concentration of 10 copies/µL (**Table 3**). The PCR performances of the two platforms are plotted and compared in **Figure 1**. Spearman’s correlation of these two assays is 0.96, which shows good consistency between the MT platform and the benchtop platform [13,14]. The paired t-test shows that there is a difference (P = 0.0002) in the Ct values between these two assays: The MT assay’s Ct value is 2 units smaller than that of the USDA benchtop assay [13,14]. These results validate the MT platform’s performance in detecting the HPAI A(H5N1) virus in raw milk.

**Table 3.** Ct values of rtPCR run of 15 raw milk samples spiked with inactivated HPAI A(H5N1) virus using the MT and the benchtop platforms. Two independent PCR runs were performed for each sample using each platform. PC: positive control. NTC: no template control. ND: not detected.

| Ct values of two PCR assays |  |  |  |  |
| --- | --- | --- | --- | --- |
| Farm ID +Virus Sample ID<br>(10 copies/μL) | Ezextractor +<br>MiniQ (MT) | | RNeasy + QS 5<br>(benchtop) | |
| Farm 1 + Sample 1 | 34.8 | 35.8 | 35.4 | 37.0 |
| Farm 2 + Sample 1 | 30.7 | 30.7 | 34.7 | 35.3 |
| Farm 3 + Sample 1 | 29.8 | 29.8 | 32.3 | 31.6 |
| Farm 4 + Sample 1 | 31.3 | 31.2 | 33.2 | 33.0 |
| Farm 5 + Sample 1 | 30.2 | 30.2 | 34.6 | 36.1 |
| Farm 6 + Sample 2 | 30.7 | 31.3 | 33.0 | 32.5 |
| Farm 7 + Sample 2 | 31.0 | 30.7 | 32.8 | 32.8 |
| Farm 8 + Sample 2 | 30.9 | 31.1 | 31.8 | 32.2 |
| Farm 9 + Sample 2 | 32.0 | 31.4 | 32.5 | 30.8 |
| Farm 10 + Sample 2 | 31.3 | 31.4 | 32.4 | 32.5 |
| Farm 11 + Sample 3 | 31.4 | 31.5 | 35.1 | 35.7 |
| Farm 12 + Sample 3 | 31.2 | 31.4 | 33.4 | 33.0 |
| Farm 13 + Sample 3 | 32.2 | 32.2 | 35.8 | 36.6 |
| Farm 14 + Sample 3 | 31.6 | 31.7 | 31.0 | 31.3 |
| Farm 15 + Sample 3 | 31.4 | 30.8 | 35.4 | 35.5 |
| PC | 24.9 | 25.1 | 24.5 | 23.7 |
| NTC | ND | ND | ND | ND |

**Figure 1.**
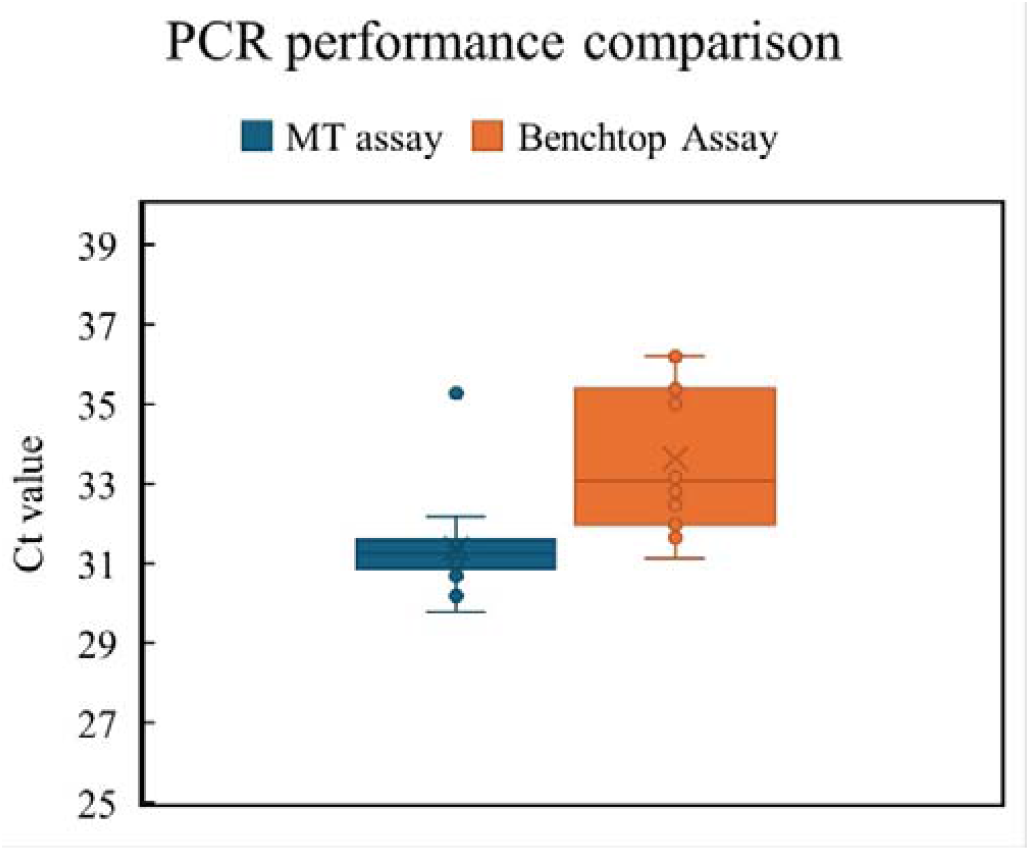
PCR performance comparison of the MT assay and the Benchtop Assay. Comparison of the performance of the MT vs. the benchtop PCR testing platforms. The Ct values listed in **Table 3** from the two platforms are analyzed. Spearman’s correlation is calculated to be 0.96, and a paired Student’s t-test is calculated to be P = 0.0002. The difference of the mean Ct values of the MT assay and the USDA-benchtop assay is calculated to be -2.3.

### PCR efficiency of the MT platform

Raw milk samples from Farm 8 and Farm 9, which underwent serial dilutions, were tested using the MT platform to determine its detection sensitivity and efficiency. The results are listed in **Table 4**. The Ct values of two independent runs of each sample at the five different dilution levels with estimated viral RNA concentrations of 1000, 100, 10, 1.0, and 0.10 copies/µL are consistent with each other and show the expected trend according to dilutions. The Ct values as a function of dilution level are plotted in **Figure 2**. PCR standard curves in **Figure 2** show that the amplification efficiencies are 93.6% and 102.5% for Farms 8 and 9, respectively. The R^2^ values (0.987 and 0.989) of both dilution series show good PCR performance of the MT platform.

**Table 4.** Ct values of rtPCR run using the MT platform of raw milk samples from two farms spiked with inactivated virus following serial ten-fold dilutions. Ct values of rtPCR run using the MT platform of raw milk samples from two farms spiked with inactivated HPAI A(H5N1) virus following serial ten-fold dilutions. Two independent PCR runs were performed for each sample. PC: positive control. NTC: no template control. ND: not detected. Note: the first (10^1^) dilution was made by adding 40 µL of inactivated virus sample to 100 µL of milk and 260 µL PBS to achieve a virus density of 1000 copies/µL. Subsequent ten-fold dilutions were made by adding 100 µL of prior dilution to 900 µL of diluent described in **Table 2**.

| Farm ID/Dilution factor | Ct values of the dilution series |  |
| --- | --- | --- |
| Farm 8/10<sup>1</sup> | 24.1 | 23.9 |
| Farm 8/10<sup>2</sup> | 26.1 | 26.1 |
| Farm 8/10<sup>3</sup> | 31.1 | 31.1 |
| Farm 8/10<sup>4</sup> | 33.7 | 34.0 |
| Farm 8/10<sup>5</sup> | 38.0 | 37.1 |
| Farm 9/10<sup>1</sup> | 23.7 | 23.6 |
| Farm 9/10<sup>2</sup> | 27.4 | 27.3 |
| Farm 9/10<sup>3</sup> | 31.3 | 31.1 |
| Farm 9/10<sup>4</sup> | 34.0 | 34.2 |
| Farm 9/10<sup>5</sup> | 36.1 | 37.0 |
| PC | 25.1 | 25.1 |
| NTC | ND | ND |

**Figure 2.**
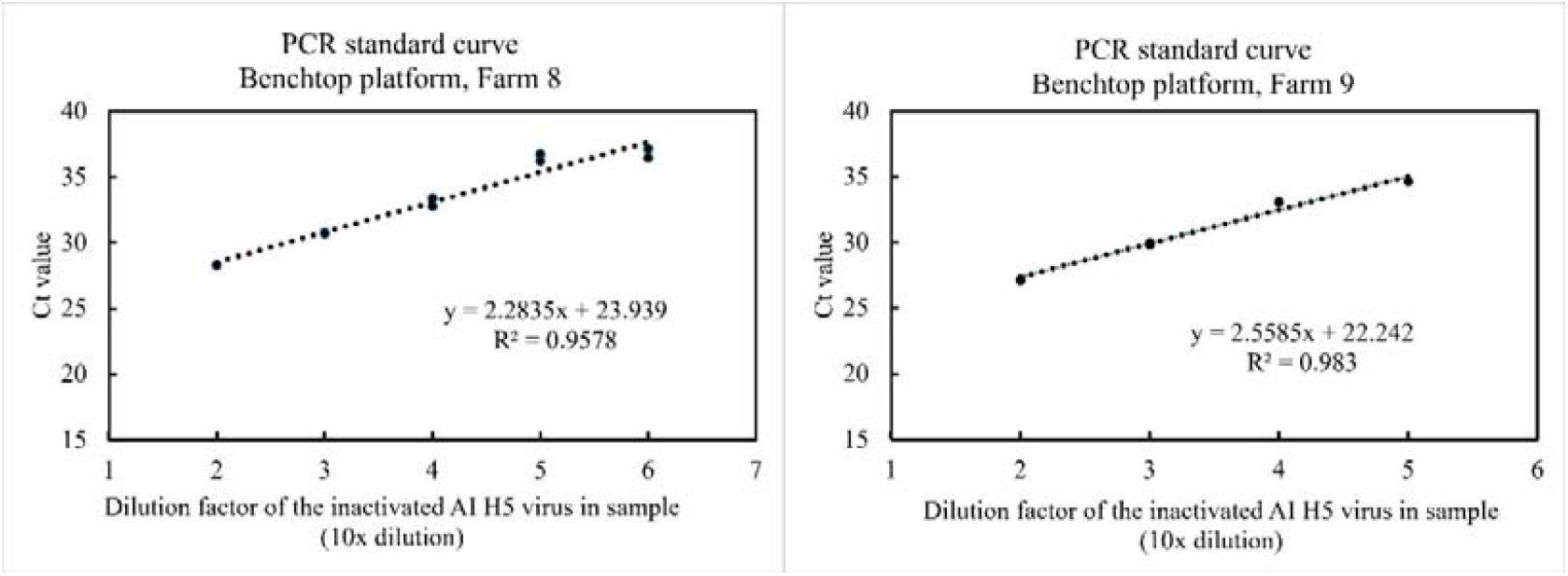
PCR standard curve of the MT platform. PCR standard curves. Ct values as a function of dilution level, from the data in **Table 4** are plotted. The amplification efficiency is calculated to be 93.6% for Farm 8 samples and 102.5% for Farm 9 samples.

**Figure 3.**
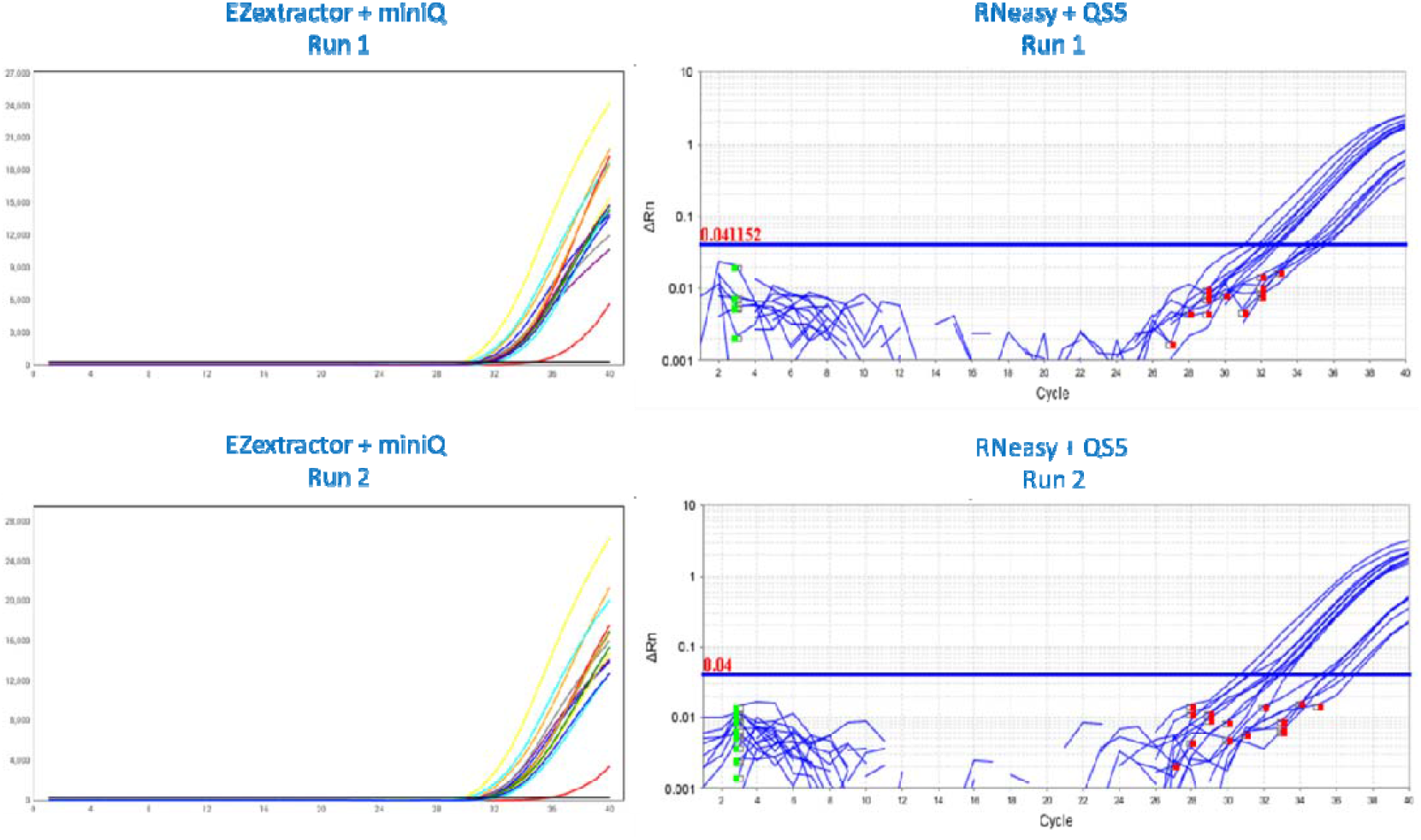
Amplification Plots of the MT assay. Amplification plots of two qPCR runs for the detection of inactivated AIV H5 spiked into raw milk samples from 15 farms. The qPCR runs were performed using the MT platform (EZextractor for RNA extraction and MiniQ thermocycler; right panels) or the benchtop platform (RNeasy for RNA extraction and QS5 thermocycler; left panels).

**Figure 4.**
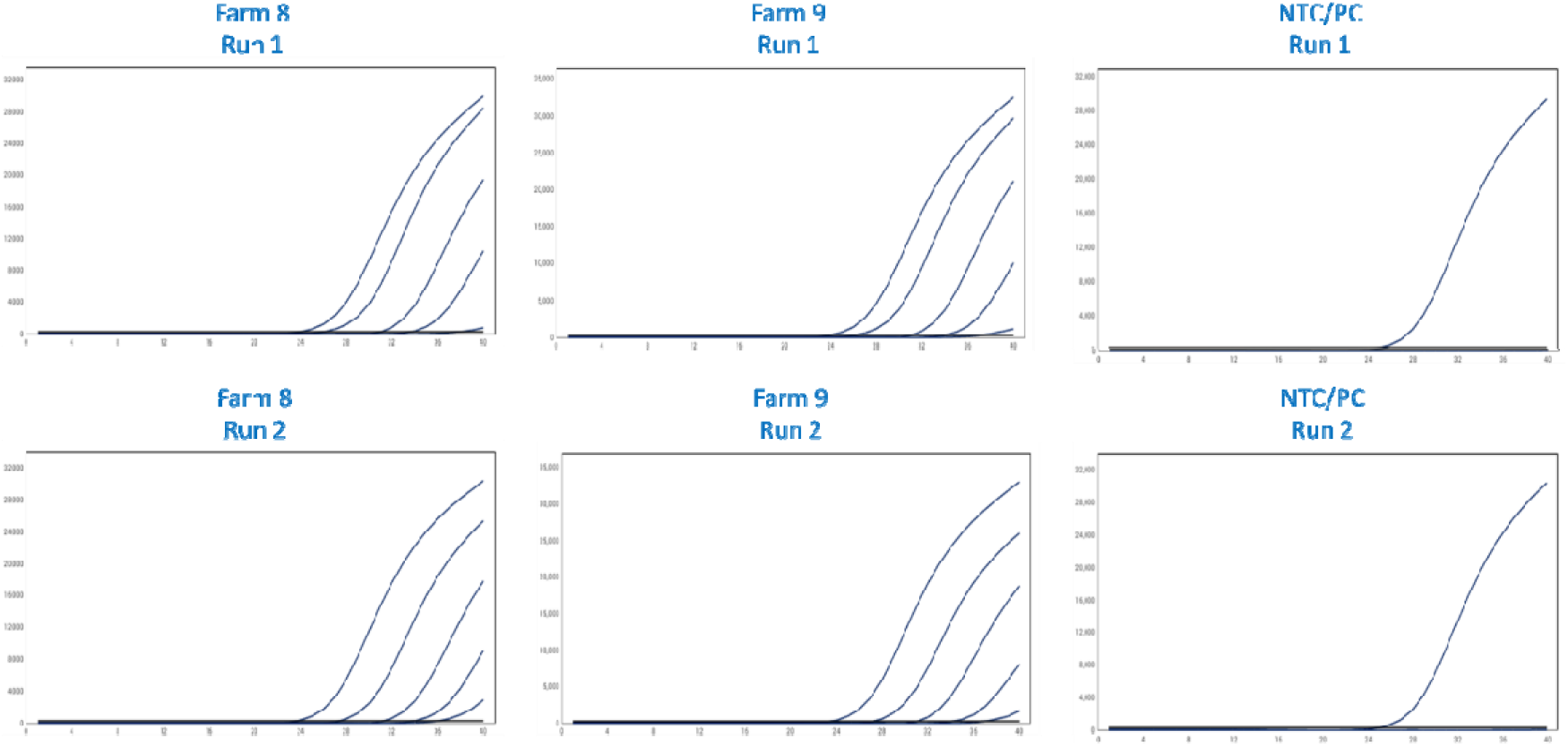
Amplification Plots of the Benchtop assay. Amplification plots of two qPCR runs for the detection of AIV H5 in serial ten-fold dilutions of inactivated AIV H5 spiked into raw milk samples from two farms. The qPCR runs were performed using the MT platform (EZextractor for RNA extraction and MiniQ thermocycler).

### Effect of Raw Milk Inhibitors on PCR Detection

The data obtained using both platforms (**Table 3**) clearly show their effectiveness in detecting inactivated HPAI A(H5N1) virus in raw milk. **Table 5** reports a comparison of diagnoses performed on raw milk vs. PBS samples spiked with the same virus density. All runs on the internal positive control (the Xeno™ RNA Control in Thermo Fisher’s AIV RNA kit) spiked samples, whether they contained raw milk or not, showed similar Ct values. In contrast, the runs on inactivated viruses showed notable differences in Ct values: The runs on samples containing raw milk showed consistently 1.5 higher Ct values than the pure PBS samples. Further, it appears that whether the combination of Ezextractor + QS5 or Ezextractor + MiniQ, the Ct values for both virus- and internal positive control-spike samples appear similar.

**Table 5.** Ct values from rtPCR conducted on two HPAI A(H5N1) virus spiked in PBS and in Raw milk samples. Ct values from rtPCR conducted on two raw milk samples compared to PBS solutions. In both comparisons, the raw milk sample and the corresponding PBS solution were spiked with the same unknown quantity of inactivated HPAI A(H5N1) virus. PC: positive control. NTC: no template control. ND: not detected.

| Ezextractor + QS5 | Ct values |  |
| --- | --- | --- |
|  | HPAI A(H5N1) Virus | Internal positive control |
| PBS | 32.8, 32.6 | 30.7, 30.5 |
| Raw milk + PBS | 34.0, 34.0 | 30.0, 29.8 |
| PC | 28.1, 28.1 | 27.9, 27.9 |
| NTC | ND | ND |
| Ezextractor + MiniQ | Ct values |  |
|  | HPAI A(H5N1) Virus | Internal positive control |
| PBS | 32.5, 32.5 | 29.5, 29.3 |
| Raw milk + PBS | 34.3, 34.0 | 30.1, 29.3 |
| PC | 27.8, 27.8 | 27.0, 26.8 |
| NTC | ND | ND |

## Discussion

Milk is a complex mixture containing fat globules, proteins, sugar, and metal ions [15]. Components in milk that may interfere with the PCR detection process include calcium ions, collagen, and myoglobin. These ions/molecules can inhibit DNA polymerase or reverse transcriptase, and *Taq* polymerase-degrading plasmin [16]. To mitigate the inhibitory effect of raw milk, we first diluted milk samples in PBS, and we were able to demonstrate the feasibility of detecting HPAI A(H5N1) virus in diluted raw milk samples using both platforms. Both rtPCR systems (MiniQ and QS5) exhibited consistent performance in detecting HPAI A(H5N1) virus in raw milk samples. Virus detection sensitivity remained robust despite the presence of PCR inhibitory substances in raw milk. The results also illustrated the impact of milk from different sources on HPAI A(H5N1) virus detection. **Table 3** shows that the Ct values from the MT platform range from 30 to 32 for all samples except the raw milk from Farm 1, which has Ct values higher than 35. The Ct values from the benchtop platform show a larger variation from 31 to 37, again with the Farm 1 sample resulting the highest Ct values.

**Table 4** shows the efficiency and the sensitivity of rtPCR detection of inactivated HPAI A(H5N1) virus in raw milk samples diluted to different virus densities. The five levels of dilution correspond to approximately 1,000, 100, 10, 1, 0.1 copies/µL. Even the lowest density HPAI A(H5N1) virus can be detected by the MT platform with high PCR efficiency. The standard curves quantitatively illustrate the precision of the MT platform detection.

From the comparison of Ct values listed in **Table 5**, it is suggested that the inhibitors do affect detection efficiency. The runs on samples containing raw milk show consistently 1.5 higher Ct values than the pure PBS samples. Furthermore, by comparing the experiments performed by using the same Ezextractor but different rtPCR (QS5 vs MiniQ), it appears that the difference in detection efficiency between the benchtop vs miniaturized instruments lies primarily with the extraction method/instrument.

Examination of the Ct values in **Table 3** also shows that the MT platform displays higher sensitivity than the benchtop platform in this study. Ct values for all 15 raw milk samples from the MT platform are on average ∼2 units lower than the ones from the benchtop platform. This difference may be attributed to the different extraction methods used in these two platforms.

The MT platform, utilizing miniaturized RNA purification and rtPCR instruments for inactivated HPAI A(H5N1) virus detection in raw milk samples represents a highly sensitive, versatile, and portable approach for managing HPAI A(H5N1) outbreaks. The successful containment and mitigation of outbreaks at the earliest instance require early detection and surveillance of the virus. Given that viral loads in milk may vary depending on infection stage and severity, the ability to detect the virus at low concentrations ensures that even minimally infected herds can be identified and managed promptly. This capability is crucial for preventing the spread of infection within and between herds. Putting into perspective the recent outbreak of H5N1 in the United States dairy cattle and the associated zoonotic risks, the implementation of the MT platform offers a practical solution for minimizing the public health and economic impacts. The portability of these systems could facilitate rapid on-site testing, reducing the delay between sample collection and results and communication with government officials. This is particularly important in high-risk areas in many states that have experienced significant outbreaks.

## Sources and manufacturers

^a^: Ezextractor Nucleic Acid Extraction System is manufactured and provided by Schweitzer Biotech Company, Taipei, Taiwan.

^b^: EZcycler Mini real-time PCR (MiniQ) system is manufactured and provided by Schweitzer Biotech Company, Taipei, Taiwan.

^c^: Ezextractor Viral DNA/RNA Extraction kit is manufactured and provided by Schweitzer Biotech Company, Taipei, Taiwan.

## Notes

### Competing Interest Statement

The authors have declared no competing interest.

### Summary of Updates

Added two authors to contributed to the analysis of the milk inhibitor data and the preparation of the manuscript related to the newly added data analysis and discussion

### Funding

- DiaVac Biotech Co.
- Schweitzer Biotech Co.

## REFERENCES

[1] Shi J, Zeng X, Cui P, Yan C, Chen H. Alarming situation of emerging H5 and H7 avian influenza and effective control strategies. Emerg Microbes Infect 2023;12. 10.1080/22221751.2022.2155072.

[2] Graziosi G, Lupini C, Catelli E, Carnaccini S. Highly Pathogenic Avian Influenza (HPAI) H5 Clade 2.3.4.4b Virus Infection in Birds and Mammals. Animals 2024;14:1372. 10.3390/ani14091372.

[3] Burrough ER, Magstadt DR, Petersen B, Timmermans SJ, Gauger PC, Zhang J, et al. Highly Pathogenic Avian Influenza A(H5N1) Clade 2.3.4.4b Virus Infection in Domestic Dairy Cattle and Cats, United States, 2024. Emerg Infect Dis 2024;30. 10.3201/eid3007.240508.

[4] Uyeki TM, Milton S, Abdul Hamid C, Reinoso Webb C, Presley SM, Shetty V, et al. Highly Pathogenic Avian Influenza A(H5N1) Virus Infection in a Dairy Farm Worker. New England Journal of Medicine 2024;390:2028–9. 10.1056/NEJMc2405371.

[5] Oguzie JU, Marushchak L V., Shittu I, Lednicky JA, Miller AL, Hao H, et al. Avian Influenza A(H5N1) Virus among Dairy Cattle, Texas, USA. Emerg Infect Dis 2024;30. 10.3201/eid3007.240717.

[6] U.S. Centers for Disease Control and Prevention. Current Situation: Bird Flu in Dairy Cows. Https://WwwCdcGov/Bird-Flu/Situation-Summary/MammalsHtml 2025.

[7] U.S. Centers for Disease Control and Prevention. Current Situation: Bird Flu in Dairy Cows. Https://WwwCdcGov/Bird-Flu/Situation-Summary/MammalsHtml 2024.

[8] Dyer O. Bird flu: Canadian teenager is critically ill with new genotype. BMJ 2024:q2529. 10.1136/bmj.q2529.

[9] U.S. Centers for Disease Control and Prevention. CDC Confirms First Severe Case of H5N1 Bird Flu in the United States. Https://WwwCdcGov/Media/Releases/2024/M1218-H5n1-FluHtml 2024.

[10] State of California US. Governor Newsom takes proactive action to strengthen robust state response to Bird Flu. Https://WwwGov.caGov/2024/12/18/Governor-Newsom-Takes-Proactive-Action-to-Strengthen-Robust-State-Response-to-Bird-Flu/ 2024.

[11] United States Department of Agriculture. USDA Announces New Federal Order, Begins National Milk Testing Strategy to Address H5N1 in Dairy Herds. Https://WwwUsdaGov/about-Usda/News/Press-Releases/2024/12/06/Usda-Announces-New-Federal-Order-Begins-National-Milk-Testing-Strategy-Address-H5n1-Dairy-Herds-0 2024.

[12] APHIS USD of A. APHIS Requirements and Recommendations for Highly Pathogenic Avian Influenza (HPAI) H5N1 Virus in Livestock For State Animal Health Officials, Accredited Veterinarians and Producers. Https://WwwAphisUsdaGov/Sites/Default/Files/Aphis-Requirements-Hpai-Livestock-Eng-SpPdf 2024.

[13] Ben Shabat M, Meir R, Haddas R, Lapin E, Shkoda I, Raibstein I, et al. Development of a real-time TaqMan RT-PCR assay for the detection of H9N2 avian influenza viruses. J Virol Methods 2010;168:72–7. 10.1016/j.jviromet.2010.04.019.

[14] Panzarin V, Marciano S, Fortin A, Brian I, D’Amico V, Gobbo F, et al. Redesign and Validation of a Real-Time RT-PCR to Improve Surveillance for Avian Influenza Viruses of the H9 Subtype. Viruses 2022;14:1263. 10.3390/v14061263.

[15] Haug A, Høstmark AT, Harstad OM. Bovine milk in human nutrition – a review. Lipids Health Dis 2007;6:25. 10.1186/1476-511X-6-25.

[16] Schrader C, Schielke A, Ellerbroek L, Johne R. PCR inhibitors - occurrence, properties and removal. J Appl Microbiol 2012;113:1014–26. 10.1111/j.1365-2672.2012.05384.x.

